# Independent Generation of Amyloid-β via Novel *APP* Transcripts

**DOI:** 10.1101/2025.05.13.651054

**Authors:** Emil K. Gustavsson, Emily Abel, Hannah Macpherson, Gunnar Brinkmalm, Diana Piotrowska, Aaron Z. Wagen, Kylie Montgomery, Claudio Villegas-Llerena, Tatiana A. Giovannucci, Rohan de Silva, Amanda Heslegrave, Nick Fox, Henrik Zetterberg, Henry Houlden, John Hardy, Selina Wray, Charles Arber, Mina Ryten

**Affiliations:** Genetics and Genomic Medicine, Great Ormond Street Institute of Child Health, UCL, London, UK; Dementia Research Centre, UCL Queen Square Institute of Neurology, UCL, London, UK; UK Dementia Research Institute at UCL, London, UK; Department of Neurodegenerative Disease, UCL Queen Square Institute of Neurology, UCL, London, UK; Department of Psychiatry and Neurochemistry, Institute of Neuroscience and Physiology, The Sahlgrenska Academy at the University of Gothenburg, Mölndal, Sweden; UK Dementia Research Institute at The University of Cambridge, Cambridge, UK; Reta Lila Weston Institute, UCL Queen Square Institute of Neurology, UCL, London, UK; Department of Clinical and Movement Neuroscience, UCL Queen Square Institute of Neurology, UCL, London, UK; Clinical Neurochemistry Laboratory, Sahlgrenska University Hospital, Mölndal, Sweden; Hong Kong Center for Neurodegenerative Diseases, Clear Water Bay, Hong Kong, China; Wisconsin Alzheimer’s Disease Research Center, University of Wisconsin School of Medicine and Public Health, University of Wisconsin-Madison, Madison, WI, USA; Department of Neuromuscular Disease, UCL Queen Square Institute of Neurology, UCL, London, UK; NIHR University College London Hospitals Biomedical Research Centre, London, UK; Institute for Advanced Study, The Hong Kong University of Science and Technology, Hong Kong SAR, China; Department of Clinical Neurosciences, School of Clinical Medicine, The University of Cambridge, Cambridge, UK; Department of Genomic Medicine, School of Clinical Medicine, The University of Cambridge, Cambridge, UK; Centro de Investigación de Genética y Biología Molecular, Instituto de Investigación, Facultad de Medicina Humana, Universidad de San Martín de Porres, Perú; School of Biology, Faculty of Health Sciences, Universidad Peruana de Ciencias Aplicadas (UPC), Lima 15023, Perú

## Abstract

The amyloid precursor protein (*APP*) is processed by multiple enzymes to generate biologically active peptides, including amyloid-β (Aβ), which aggregates to form the hallmark pathology of Alzheimer’s disease (AD). Aβ is produced through an initial β-secretase cleavage of APP, generating a 99-amino acid C-terminal fragment (APP-C99). Subsequent cleavage of APP-C99 by γ-secretase produces Aβ peptides of varying lengths. To better understand the transcriptional regulation of Aβ production, we employed long-read RNA sequencing and identified previously unannotated transcripts encoding APP-C99 with an additional methionine residue (APP-C100), generated independently of β-secretase cleavage. These transcripts are expressed separately from full-length *APP*, and we observed that cells lacking full-length *APP* can still produce Aβ through these shorter isoforms. Importantly, mass spectrometry analysis of cerebrospinal fluid (CSF) revealed peptides consistent with the methionine-extended Aβ species, supporting the *in vivo* translation of these transcripts. Our findings reveal an alternative pathway for Aβ generation and aggregation, highlighting a potential new target for modulating Aβ accumulation in AD.

---

Alzheimer’s disease (AD) is the primary cause of dementia, the most common neurodegenerative disorder, and ranks as the fifth leading cause of death globally (*1*). The main pathological hallmark of AD is the formation of amyloid fibrils that accumulate into plaques, and a progressive loss of neuronal cells (*2*). These plaques are primarily composed of amyloid-β (Aβ) peptides, which are produced through the proteolytic cleavage of the amyloid precursor protein (APP) (*2–5*).

Beyond Aβ, the amyloid precursor protein (APP) is processed by multiple enzymes, including α-, β-, γ-, and η-secretases, generating a variety of biologically active protein fragments (*6*). Some of these fragments function in the extracellular space to regulate synaptic activity, such as soluble APPα (sAPP⍺) and soluble APPη (sAPPη) (*7*, *8*). Others, like the APP intracellular domain (AICD), are implicated in transcriptional regulation (*9*). Additionally, membrane-bound peptides such as C99 have been implicated in autophagy (*10*).

In the amyloidogenic pathway, APP is initially cleaved by β-secretase (β-APP-cleaving enzyme-1, BACE1) within its extracellular domain, producing a large soluble fragment known as soluble peptide APPβ (sAPPβ) and a smaller, membrane-bound fragment called C99 (or β-CTF) (*4*, *11–13*). C99 is then further processed by γ-secretase, a multi-subunit enzyme complex that cleaves within APP’s transmembrane region, releasing amyloid-β (Aβ) peptides of varying lengths, typically ranging from 37 to 43 residues (*14*). Among these, Aβ42 and Aβ43 are particularly prone to aggregation due to their hydrophobic nature and tendency to form β-sheet structures, making them a major contributor to the formation of amyloid plaques.

Over the past three decades, the amyloid hypothesis has been extensively tested, making amyloid a key therapeutic target for AD (*12*, *15*). As a result, three anti-amyloid monoclonal antibodies (mAbs) designed to reduce brain Aβ have received FDA approval: Aducanumab (Aduhelm; Biogen, Cambridge, MA, USA) (*16*), Lecanemab (Leqembi; Eisai Inc. and Biogen, Cambridge, MA, USA) (*17*, *18*), and Donanemab (Kisunla; Eli Lilly, Indianapolis, IN, USA) (*19*, *20*). The observed slowing of cognitive decline in early disease stages with these treatments reinforces the central role of Aβ in AD.

APP is encoded by the *APP* gene on chromosome 21 where the 18 currently annotated exons (GENCODE46) generate 20 transcripts that are primarily translated into three major isoforms: APP-695, APP-751, and APP-770 (*21–23*). Quantitative changes in *APP* expression due to multiplications of the locus lead to cognitive impairment associated with AD biological signatures in patients with Down syndrome (*24*, *25*), and is associated with AD-like dementia (*26*, *27*). Similarly, qualitative changes in *APP* processing due to missense mutations either in the substrate (*APP*) or active component of γ-secretase (*PSEN1*, *PSEN2*) cause dominant, early-onset forms of AD (*28*).

Despite these well-established links, the full extent of *APP* transcriptional diversity has received little attention. To address this, we conducted targeted high-depth long-read sequencing of the *APP* locus to comprehensively characterize its transcript diversity. We further assessed the coding potential of all *APP* transcripts using RNA from brain tissue and brain-relevant induced pluripotent stem cells (iPSC)-derived cell types.

## Results

### Long-read RNA-seq reveals unannotated transcripts of *APP* in the human brain and brain-relevant cell types

Annotation inaccuracies are well-documented, with long-read sequencing revealing numerous novel transcripts, including previously uncharacterized protein isoforms both across the transcriptome (*29–32*), and within well-studied disease-associated genes (*33*, *34*). To gain deeper insight into *APP* transcription, we performed targeted long-read RNA sequencing across a range of cell types, including iPSC, neuroepithelial cells, neural progenitor cells, and iPSC-derived cortical neurons, astrocytes, and microglia, as well as post-mortem brain samples (**Fig. 1A**; **table S1**).

**Fig. 1.**
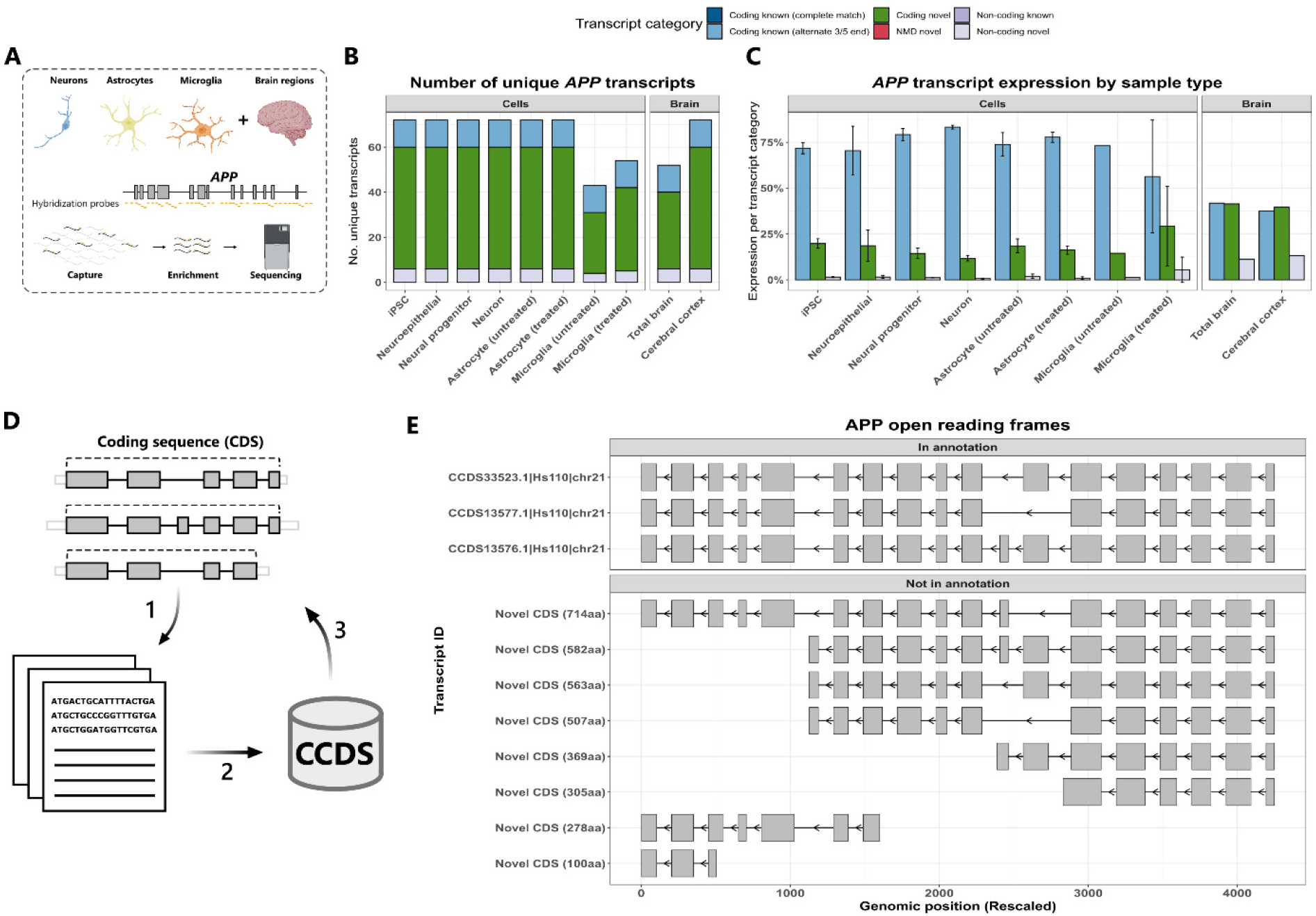
Targeted long-read RNA-seq of *APP* identifies frequent novel transcription. **(A)** Schematic outlining targeted long-read RNA-seq paradigm. **(B)** Bar chart depicting the number of unique *APP* transcripts identified per transcript category through targeted long-read RNA-seq in iPSC, neuroepithelial cells, neural progenitor cells, iPSC-derived cortical neurons, astrocytes, microglia, and in brain tissue. **(C)** Stacked bar chart showing expression per transcript category of APP in iPSC, neuroepithelial cells, neural progenitor cells, iPSC-derived cortical neurons, astrocytes, microglia, and in brain tissue. **(D)** Schematic outlining the analysis of unique Coding DNA Sequences (CDS) across transcripts. In step one reading frames are predicted to retain CDS for each transcript. In step two, the CDS are all compared to NCBI CCDS database. In step three, the CDS of APP transcripts are annotated with this information to know whether the CDS is novel or in annotation. **(E)** Visualisation of 11 unique *APP* CDS sequences identified.

We employed *APP*-targeted PacBio Iso-Seq across 26 human samples (**table S1**), whose circular consensus sequencing enabled >99% base pair accuracy, ensuring precise mapping. Stringent quality control and collapsing mapped reads resulted in the identification of 1,626 unique *APP* transcripts, each supported by ≥2 full-length HiFi reads across all samples (**Fig. S1**). To prioritise consistently expressed transcripts, we applied a threshold requiring each transcript to be detected in at least 80% of samples, with a minimum relative expression of 0.1% per sample. This corresponded to an average of 21 to 3,971 full-length HiFi reads per transcript, resulting in 72 unique *APP* transcripts that met these criteria (**Fig. 1B; table S2**).

We predicted open reading frames (ORFs) for all transcripts using two approaches: SQANTI3 (34), which applies the GeneMarkS-T (GMST) algorithm (*35*) to identify ORFs from transcript sequences, and ORFik (*36*). Among the 72 identified transcripts, 54 were predicted to be translated, evade nonsense-mediated decay (NMD), and lack a full splice match with the reference, classifying them as novel coding transcripts. (**Fig. S2**). A further twelve transcripts were predicted to be stable, translated, and contained exon-exon junctions that fully matched with the reference but exhibited alternative 3′ and/or 5′ ends so classifying them as known coding transcripts (alternate 3/5 end) (**Fig. S3**). The remaining six transcripts were predicted to be non-coding and lacked a full splice match with the reference, classifying them as non-coding novel transcripts (**Fig. S4**).

The largest proportion of *APP* expression came from known coding (alternate 3/5 end) transcripts, with a mean relative expression of 73.2 ± 8.0% in cells and 39.7 ± 3.1% in brain tissue. This was followed by novel coding transcripts, with mean relative expression of 17.9 ± 5.3% in cells and 40.6 ± 1.2% in brain tissue. Non-coding novel transcripts had the lowest expression, with mean relative expression of 1.8 ± 1.6% in cells and 12.2 ± 1.3% in brain tissue (**Fig. 1C**).

By collapsing predicted coding transcripts based on their coding DNA sequence (CDS) (**Fig. 1D**), we identified 11 unique open reading frames (ORFs). Three of these ORFs corresponded to annotations in the NCBI Consensus CDS (CCDS) database (PMID: 19498102), while the remaining seven were novel (**Fig. 1E**).

### *APP* transcripts are predicted to encode cleavage products of APP

In the amyloidogenic pathway, β-secretase cleaves APP within its extracellular domain at KM|D (APP695: 671–672), generating soluble APPβ (sAPPβ) and the membrane-bound C99. Notably, when comparing start codons (ATG) of the novel CDS and their resulting peptide sequences to known secretase cleavage sites one ORF overlaps with a β-secretase cleavage site. This ORF (Novel 100aa) is derived from two transcripts, PB.4.557 and PB.4.251 (**Fig. 2A**). Since usage of unannotated 5′ transcription start sites (TSSs) was a common feature of these two *APP* transcripts (**Fig. 2A**), we focused on validating these sites using cap analysis gene expression (CAGE) peaks (defined by FANTOM5)(*37*, *38*). We found that, even though CAGE sequencing only captures the first 20 to 30 nucleotides from the 5′-end, these novel *APP* 5′ TSSs were located within 50 bp of CAGE peaks, providing additional confidence in the calling of these transcripts (**Fig. 2B**). Furthermore, we noted that already in annotation there is a transcript (ENST00000464867) classified as non-coding which closely matches both PB.4.557 and PB.4.251, albeit with a slightly different TSS and TTS (**Fig. S5**).

**Fig. 2.**
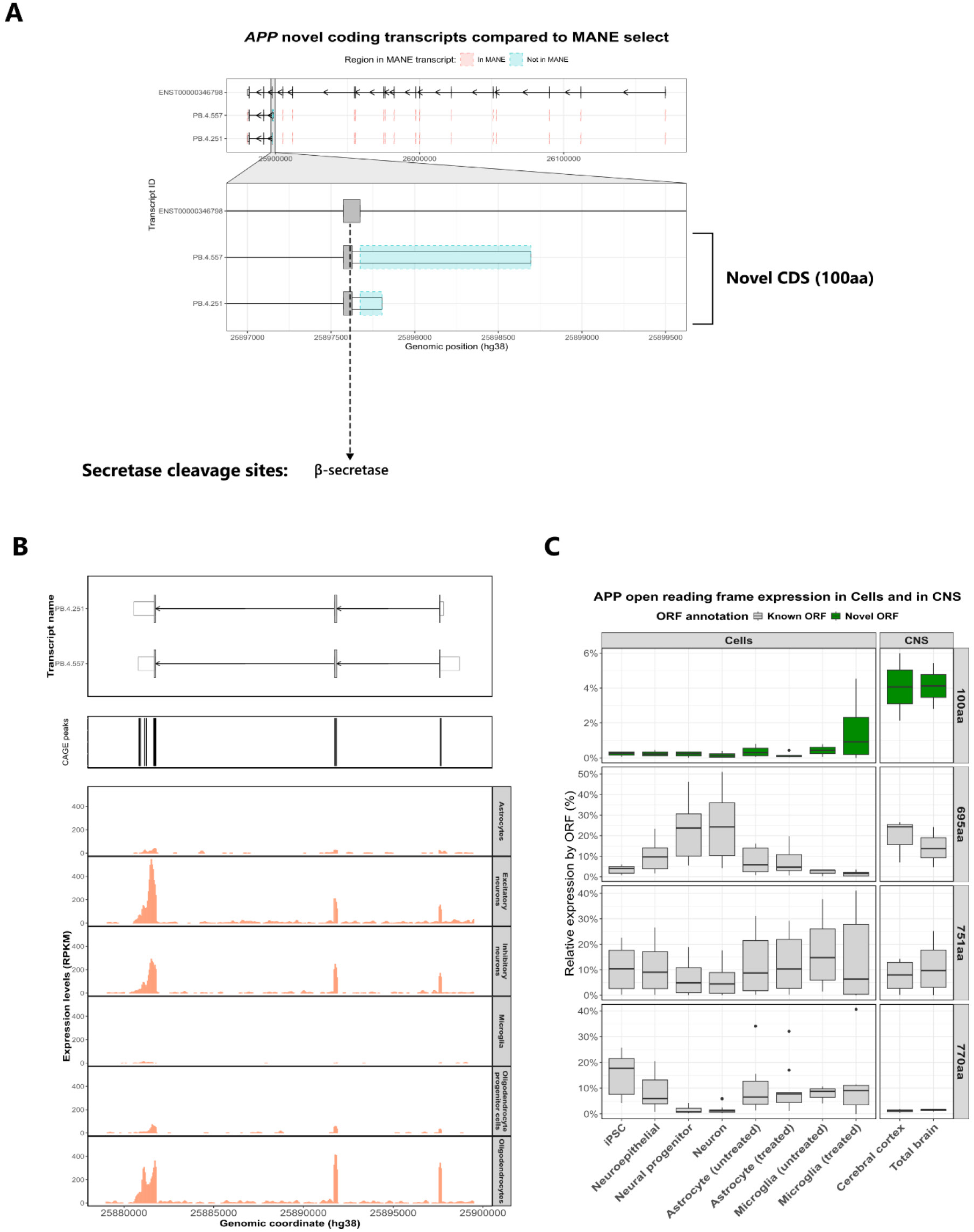
APP cleavage peptides can be generated independently from full length APP via novel transcriptional events. **(A)** Novel coding APP transcripts are visualised using ggtranscript. Differences compared to the MANE Select transcript (ENST00000346798) are shown in blue (novel regions) and red (missing regions). Two transcripts (PB.4.557 and PB.4.251) are predicted to encode a 100-amino-acid peptide (Novel CDS (100aa)) corresponding to the 99-amino-acid C-terminal fragment (APP-C99), with an added methionine, independent of β-secretase cleavage. **(B)** Novel protein coding *APP* transcript on top aligned with 5′ snRNA-seq expression in human dorsolateral prefrontal cortex (DLPFC) and CAGE sequencing data from FANTOM5. **(C)** Relative expression of these novel open reading frames compared to annotated ones in brain-relevant iPSC-derived cell types and across post-mortem brain regions (“treated” refers to cultures exposed to inflammatory cues; lipopolysaccharide in microglia and TNFα/IL1α/C1q in astrocytes).

We then used 5′ single-nucleus RNA-seq (snRNA-seq) of the human dorsolateral prefrontal cortex (DLPFC) to assess the expression of *APP* across various cell types, including astrocytes, excitatory neurons, inhibitory neurons, microglia, oligodendrocytes, and oligodendrocyte precursor cells (OPCs) (**Fig. 2B**). Our analysis revealed the strongest signal for the ORF (Novel 100aa) in oligodendrocytes, followed by excitatory and inhibitory neurons. In contrast, we observed an absence of signal at the first exon of PB.845.2888 (APP) in microglia, along with an overall lower expression of novel APP transcripts in microglia and OPCs (**Fig. 2C**).

### Translation of APP C100 in human cerebrospinal fluid

This ORF (Novel 100aa) is generated from two transcripts (PB.4.557 and PB.4.251) that are expected to produce the β-secretase cleavage product APP-C99, with an additional N-terminal methionine, hereafter known as C100 (**Fig. 3A**). Our mass spectrometry analysis of cerebrospinal fluid (CSF) provided strong evidence for the translation of C100 (**Fig. 3B**). Specifically, we identified three peptides: Aβ -1 to 15, Aβ -1 to 15 oxidised at M(-1), and Aβ -1 to 16, with excellent sequence coverage, leaving no doubt about their identity. The -1 position refers to the methionine located one amino acid upstream of the β-secretase cleavage site. To enhance detection, we processed 5 mL of CSF, first depleting 1-x and x-40 peptides—though this step was only partially successful, followed by immunoprecipitation with 6E10 and 4G8 antibodies. While the number of MS/MS spectra obtained for these peptides was relatively low compared with others, such as Aβ -3 to 15, Aβ -2 to 15, and particularly Aβ 1-15, these findings provide critical support for the translation of novel *APP* transcripts, and reinforce the validity of our transcriptomic discoveries at the proteomic level.

**Fig. 3:**
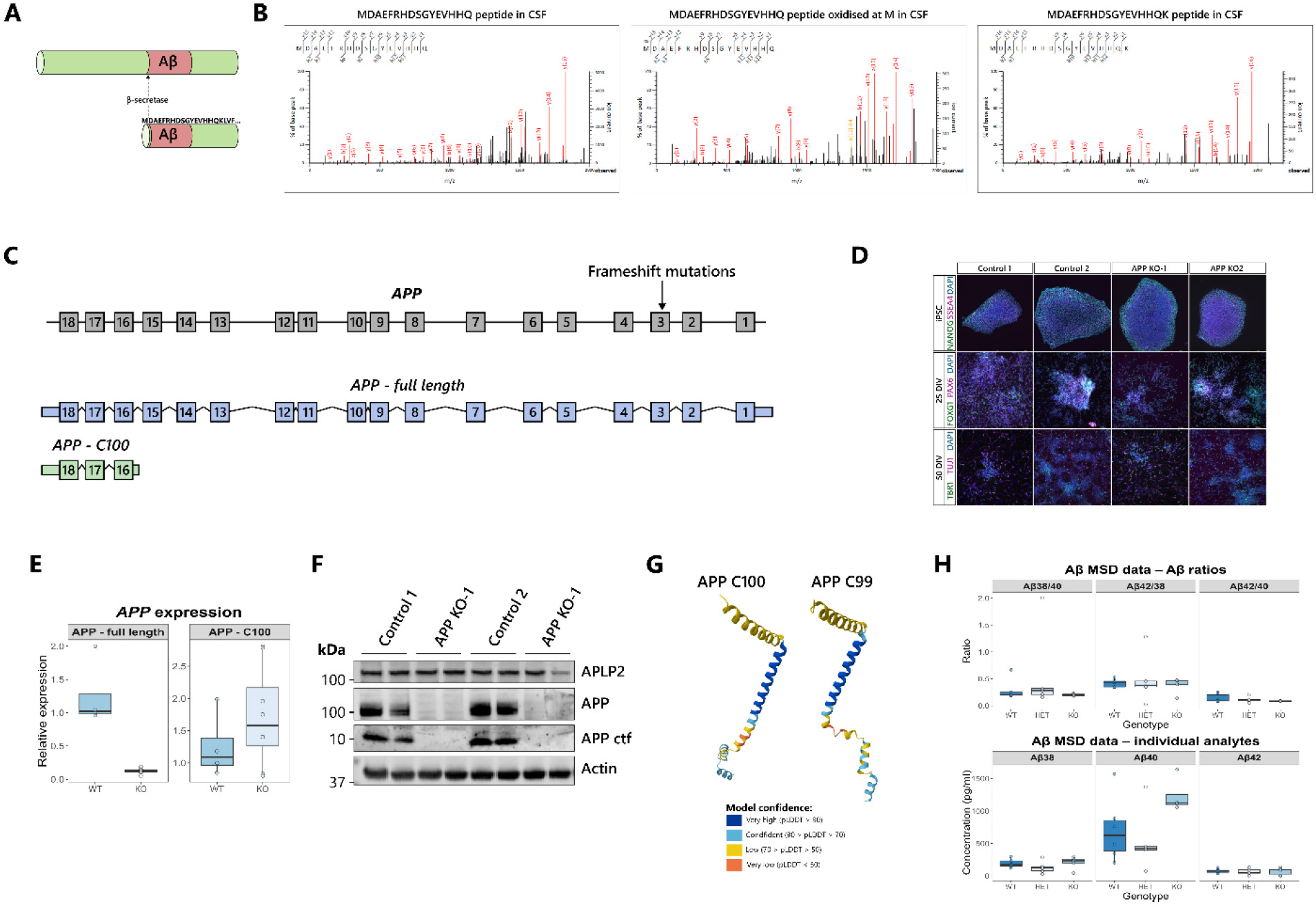
Generation of Aβ from APP-C100. **(A)** Schematic showing the predicted peptide translated from APP-C100. **(B)** Top-down mass spectrometry in human cerebrospinal fluid (CSF) identifies three independent peptides that include a Methionine as an N terminal extension to Aβ, supporting translation from APP-C100 rather than β-secretase cleavage. **(C)** Strategy for generation of APP full-length KO cells via the introduction of frameshift mutations in exon 3 of APP. This strategy leaves APP-C100 unaffected. **(D)** Immunocytochemical characterisation of APP full-length KO iPSCs, whereby KO cells exhibit characteristic pluripotency marker expression (OCT4 and SSEA4) top, markers of cortical neuron patterning at the precursor stage (25 DIV, FOXG1 and PAX6) middle, and deep layer cortical glutamatergic neuronal marker TBR1 with pan neuronal marker TUJ1, at 50 DIV bottom. **(E)** qPCR analysis of APP and APP-C100 expression in unedited iPSCs or APP full-length KO iPSCs (n=6 for each of one control clone and two APP full-length KO clonal iPSC lines). **(F)** Western blotting of APP, APP C-terminal fragment, and APLP2 in iPSC-derived cortical neurons (100 DIV). **(G)** AlphaFold3 predictions of APP-C100 and APP-C99. **(H)** Electrochemiluminescent analysis for Aβ peptides (Aβ38, Aβ40 and Aβ42) in iPSC-derived neuronal conditioned media. Data presented as ratios, or normalised to the protein content of the pellet from the corresponding wells (n=7; 3-4 independent inductions for 2 control clones and 2 APP full-length KO clones each).

### Novel *APP* transcripts generate amyloid-β peptides independently of full-length *APP* transcription

To determine whether novel *APP*-C100 transcripts could be generated independently of *APP* transcripts containing full-length ORFs, we used CRISPR-Cas9 genome editing to introduce frameshift mutations in exon 3 of the *APP* gene, generating two independent full-length APP knockout (KO) iPSC clones (**Fig. 3C-D**). This approach disrupted full-length *APP* transcription, rendering *APP* mRNA undetectable by qPCR and full-length APP protein undetectable by Western blotting in iPSC-derived neuronal cultures (**Fig. 3E-F**). Despite this, expression of the novel *APP*-C100 transcripts in full-length APP KO iPSCs remained comparable to unedited controls (**Fig. 3E-F**).

The predicted structure of APP-C100 closely resembles that of APP-C99 (**Fig. 3G**), suggesting that APP-C100 may also contribute to Aβ generation. To test whether APP-C100 can produce Aβ independently of full-length APP, we differentiated both full-length APP KO and unedited iPSCs into cortical neurons (**Fig. 3D**). Surprisingly, Aβ peptides were detectable in APP KO neurons, with ratios indistinguishable from unedited neurons, despite the absence of full-length APP protein (**Fig. 3H**). Aβ1-40 and Aβ11-40 were also detectable in full-length APP KO neurons by IP-mass spectrometry, albeit 10-fold lower than unedited controls. These findings suggest that neurons lacking full-length *APP* transcription remain capable of generating Aβ in iPSC-derived cultures.

## Discussion

Traditionally, Aβ peptides have been thought to arise exclusively from the sequential cleavage of full-length APP. However, our findings indicate an additional transcriptional mechanism that generates cleavage products independently of full-length APP production.

APP undergoes extensive proteolytic processing, generating multiple fragments with diverse biological functions. Therefore, transcript diversity may have arisen to enable the generation of the same cleavage products in a manner that was distinct and independent of other peptides domains.

The discovery of APP-C100 transcripts that can generate Aβ independently of BACE1 cleavage, presents a significant challenge to the efficacy of BACE1 inhibitors in AD treatment. This alternative pathway suggests that Aβ production can continue even when BACE1 activity is inhibited, potentially explaining the lack of clinical success of BACE1 inhibitors, which showed only partial reduction in amyloid levels (*39*). This residual Aβ production highlights the need for combination therapies targeting multiple steps in the amyloidogenic pathway, such as incorporating γ-secretase modulators or stabilizers, to address both BACE1-dependent and independent Aβ generation. Consequently, a more comprehensive approach that tackles various aspects of Aβ production may be necessary to achieve more effective therapeutic outcomes for AD patients.

The C99 fragment of APP has been extensively studied in isolation from full length APP. *In vitro* studies have employed the overexpression of C99 to study the effect of familial Alzheimer’s disease (fAD) mutations in *PSEN1*, whereby the efficiency of cleavage of C99 and the subsequent processing of Aβ correlates with the age of onset for each mutation (*40*, *41*). Indeed, overexpression of the C99 fragment alone recapitulates AD disease phenotypes in rodent models, such that Aβ deposition, plaque formation and neurodegeneration was observed (*42*). Furthermore, C99 has been shown to accumulate within neurons as an early, disease-associated phenotype (*43*). Together these studies independently validate the biological relevance of the novel transcript presented here.

It is also important to consider fAD mutations in the setting of these novel transcripts. To date, 114 mutations have been found in *APP* that lead to fAD (Alzforum). It is striking that every mutation that is classed as “pathogenic” or “likely pathogenic” lies at codon 670 or more C-terminal, meaning that fAD mutations are wholly enriched within the APP-C100 transcript.

Finally, it is intriguing to consider the relative abundance of the APP cleavage products. In healthy control CSF, sAPPα is found at 200-400ng/ml whereas Aβ is around 1-2ng/ml (over 20-fold difference in abundance, accounting for the molecular weight (*44*)). These concentrations support the notion that Aβ is a minor product of APP processing and opens the possibility that Aβ may be generated from rare transcripts.

In summary, we establish that Aβ can be generated from transcriptional events that are independent of canonical *APP* transcription. This enables the generation of biologically active peptides in an energetically efficient manner, distinct from other cleavage products. This process is not universal (we did not observe similar transcripts in *APP* family members *APLP1* and *APLP2*, nor other dementia associated genes that undergo complex cleavage events such as *GRN*), further emphasising the biological relevance of this transcriptional event for APP physiology. Therefore, these transcripts offer novel targets for lowering AD pathogenic peptides with reduced off-target considerations.

## ONLINE METHODS

## PACBIO TARGETED ISO-SEQ

### Cell Culture

#### iPSC, neuroepithelial, neural progenitor, cortical neuron, astrocyte, and microglia cells

Control iPSCs consisted of the previously characterized lines Ctrl1 (*45*), ND41866 (Coriel), RBi001 (EBiSC/Sigma) and SIGi1001 (EBiSC/Sigma) as well as the isogenic line previously generated (*46*). Reagents were purchased from Thermo Fisher Scientific unless otherwise stated. iPSCs lines were grown in Essential 8 media on geltrex substrate and passaged using 0.5mM EDTA. Cortical neurons were differentiated using dual SMAD inhibition for 10 days (10µM SB431542 and 1µM dorsomorphin, Tocris) in N2B27 media before maturation in N2B27 alone (*47*). Day 100 ± 5 days was taken as the final timepoint, with day 10 as neuroepithelial, and day 40 as neural progenitors. Astrocytes were generated following a similar neural induction protocol until day 80 before repeatedly passaging cortical neuronal inductions in 10ng/ml FGF2 (Peprotech) to enrich for astrocyte precursors. At day 150, to generate mature astrocytes, a two-week maturation consisted of BMP4 (10ng/ml, Thermo Fisher) and LIF (10ng/ml, Sigma) (*48*). To induce inflammatory conditions, astrocytes were stimulated with TNFα (30ng/ml, Peprotech), IL1α (3ng/ml, Peprotech) and C1q (400ng/ml, Merck) (*49*). iPSC-microglia were differentiated following the protocol of Xiang et al (*50*). Embryoid bodies were generated using 10,000 iPSCs and myeloid differentiation was initiated in Lonza XVivo 15 media, IL3 (25ng/ml, Peprotech) and MCSF (100ng/ml, Peprotech). Microglia released from embryoid bodies were harvested weekly from 4 weeks and matured in DMEM-F12 supplemented with 2X insulin/transferrin/selenium, 1X N2 supplement, 1X glutamax, 1X NEAA and 5ng/ml insulin supplemented with IL34 (100ng/ml, Peprotech), MCSF (25ng/ml, Peprotech), TGFβ1 (5ng/ml, Peprotech). A final two-day maturation consisted of CXC3L1 (100ng/ml, Peprotech) and CD200 (100ng/ml, 2B Scientific). Inflammation was stimulated with lipopolysaccharide (100ng/ml, Sigma). Total RNA was extracted using the Qiagen RNeasy kit according to the manufacturer’s protocol with β-mercaptoethanol added to buffer RLT and with a DNase digestion step included.

### cDNA synthesis

A total of 250ng of RNA was used per sample for reverse transcription. Two different cDNA synthesis approaches were used: (i) Human brain cDNA was generated by SMARTer PCR cDNA synthesis (Takara) and (ii) iPSC derived cell lines were generated using NEBNext® Single Cell/Low Input cDNA Synthesis & Amplification Module (New England Biolabs). For both reactions sample-specific barcoded oligo dT (12 µM) with PacBio 16mer barcode sequences were added (**Supplementary Table 3**).

#### SMARTer PCR cDNA synthesis

First strand synthesis was performed as per manufacturer instructions, using sample-specific barcoded primers instead of the 3’ SMART CDS Primer II A. We used a 90 min incubation to generate full-length cDNAs. cDNA amplification was performed using a single primer (5’ PCR Primer II A from the SMARTer kit, 5′ AAG CAG TGG TAT CAA CGC AGA GTA C 3′) and was used for all PCR reactions post reverse transcription. We followed the manufacturer’s protocol with our determined optimal number of 18 cycles for amplification; this was used for all samples. We used a 6 min extension time in order to capture longer cDNA transcripts. PCR products were purified separately with 1X ProNex® Beads.

#### NEBNext® Single Cell/Low Input cDNA Synthesis & Amplification Module

A reaction mix of 5.4 μL of total RNA (250 ng in total), 2 μL of barcoded primer, 1.6 μL of dNTP (25 mM) held at 70°C for 5 min. This reaction mix was then combined with 5 μL of NEBNext Single Cell RT Buffer, 3 μL of nuclease-free H_2_O and 2 μL NEBNext Single Cell RT Enzyme Mix. The reverse transcription mix was then placed in a thermocycler at 42°C with the lid at 52°C for 75 minutes then held at 4°C. On ice, we added 1 μL of Iso-Seq Express Template Switching Oligo and then placed the reaction mix in a thermocycler at 42°C with the lid at 52°C for 15 minutes. We then added 30 μL elution buffer (EB) to the 20 μL Reverse Transcription and Template Switching reaction (for a total of 50 μL), which was then purified with 1X ProNex® Beads and eluted in 46 μL of EB. cDNA amplification was performed by combining the eluted Reverse Transcription and Template Switching reaction with 50 μL of NEBNext Single Cell cDNA PCR Master Mix, 2 μL of NEBNext Single Cell cDNA PCR Primer, 2 μL of Iso-Seq Express cDNA PCR Primer and 0.5 μL of NEBNext Cell Lysis Buffer.

### cDNA Capture Using IDT Xgen® Lockdown® Probes

We used the xGen Hyb Panel Design Tool (https://eu.idtdna.com/site/order/designtool/index/XGENDESIGN) to design non-overlapping 120-mer hybridization probes against *APP*. We removed any overlapping probes with repetitive sequences (repeatmasker) and to reduce the density of probes mapping to intronic regions 0.2, which means 1 probe per 1.2kb. In the end, our probe pool consisted of XX probes were targeted towards *APP*.

We pooled an equal mass of barcoded cDNA for a total of 500 ng per capture reaction. Pooled cDNA was combined with 7.5 μL of Cot DNA in a 1.5 mL LoBind tube. We then added 1.8X of ProNex beads to the cDNA pool with Cot DNA, gently mixed the reaction mix 10 times (using a pipette) and incubated for 10 min at room temperature. After two washes with 200 μL of freshly prepared 80% ethanol, we removed any residual ethanol and immediately added 19 μL hybridization mix consisting of: 9.5 μL of 2X Hybridization Buffer, 3 μL of Hybridization Buffer Enhancer, 1 μL of xGen Asym TSO block (25 nmole), 1 μL of polyT block (25 nmole) and 4.5 μL of 1X xGen Lockdown Probe pool.

The PacBio targeted Iso-Seq protocol is described in detail at protocols.io (dx.doi.org/10.17504/protocols.io.n92ld9wy9g5b/v1).

### Automated Analysis of Iso-Seq data using Snakemake

For the analysis of targeted PacBio Iso-Seq data, we created two Snakemake (*51*) (v 5.32.2; RRID:SCR_003475) pipelines to analyse targeted long-read RNA-seq robustly and systematically:

**APTARS** (Analysis of PacBio TARgeted Sequencing, https://github.com/sid-sethi/APTARS): For each SMRT cell, two files were required for processing: (i) a subreads.bam and (ii) a FASTA file with primer sequences, including barcode sequences.

Each sequencing run was processed by ccs (v 5.0.0; RRID:SCR_021174; https://ccs.how/), which combines multiple subreads of the same SMRTbell molecule and to produce one highly accurate consensus sequence, also called a HiFi read (≥ Q20). We used the following parameters: --minLength 10 –maxLength 50000 – minPasses 3 –minSnr 2.5 –maxPoaCoverage 0 –minPredictedAccuracy 0.99.

Identification of barcodes, demultiplexing and removal of primers was then performed using lima (v 2.0.0; https://lima.how/) invoking –isoseq –peek-guess.

Isoseq3 (v 3.4.0; https://github.com/PacificBiosciences/IsoSeq) was then used to (i) remove polyA tails and (ii) identify and remove concatemers using, with the following parameters refine –require-polya, --log-level DEBUG. This was followed by clustering and polishing with the following parameters using: cluster flnc.fofn clustered.bam – verbose –use-qvs.

Reads with predicted accuracy ≥ 0.99 were aligned to the GRCh38 reference genome using minimap2 (*52*) (v 2.17; RRID:SCR_018550) using -ax splice:hq -uf – secondary=no. samtools (*53*) (RRID:SCR_002105; http://www.htslib.org/) was then used to sort and filter the output SAM for the locus of gene of interest, as defined in the config.yml.

We used cDNA_Cupcake (v 22.0.0; https://github.com/Magdoll/cDNA_Cupcake) to: (i) collapse redundant transcripts, using collapse_isoforms_by_sam.py (--dun-merge-5-shorter) and (ii) obtain read counts per sample, using get_abundance_post_collapse.py followed by demux_isoseq_with_genome.py.

Isoforms detected were characterized and classified using SQANTI3 (*54*) (v 4.2; https://github.com/ConesaLab/SQANTI3) in combination with GENCODE (v 38) comprehensive gene annotation. An isoform was classified as full splice match (FSM) if it aligned with reference genome with the same splice junctions and contained the same number of exons, incomplete splice match (ISM) if it contained fewer 5′ exons than reference genome, novel in catalog (NIC) if it is a novel isoform containing a combination of known donor or acceptor sites, or novel not in catalog (NNC) if it is a novel isoform with at least one novel donor or acceptor site.

PSQAN (Post Sqanti QC Analysis, https://github.com/sid-sethi/PSQAN) Following transcript characterisation from SQANTI3, we applied a set of filtering criteria to remove potential genomic contamination and rare PCR artifacts. We removed an isoform if: (1) the percent of genomic “A” s in the downstream 20 bp window was more than 80% (“perc_A_downstream_TTS” > 80); (2) one of the junctions was predicted to be template switching artifact (“RTS_stage” = TRUE); or (3) it was not associated with the gene of interest. Using SQANTI’s output of ORF prediction, NMD prediction and structural categorisation based on comparison with the reference annotation (GENCODE), we grouped the identified isoforms into the following categories: (1) Non-coding novel – if predicted to be non-coding and not a full-splice match with the reference; (2) Non-coding known – if predicted to be non-coding and a full-splice match with the reference; (3) NMD novel – if predicted to be coding & NMD, and not a full-splice match with the reference; (4) NMD known – if predicted to be coding & NMD, and a full-splice match with the reference; (5) Coding novel – if predicted to be coding & not NMD, and not a full-splice match with the reference; (6) Coding known (complete match) – if predicted to be coding & not NMD, and a full-splice & UTR match with the reference; and (7) Coding known (alternate 3’/5’ end) – if predicted to be coding & not NMD, and a full-splice match with the reference but with an alternate 3’ end, 5’ end or both 3’ and 5’ end.

Given a transcript *T* in sample *i* with *FLR* as the number of full-length reads mapped to the transcript *T*, we calculated the normalised full-length reads (*NFLR*_*Ti*_) as the percentage of total transcription in the sample:

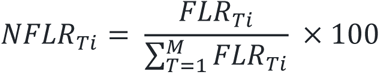

where, *NFLR*_*Ti*_ represents the normalised full-length read count of transcript *T* in sample *i*, *FLR*_*Ti*_ is the full-length read count of transcript *T* in sample *i* and *M* is the total number of transcripts identified to be associated with the gene after filtering. Finally, to summarise the expression of a transcript associated with a gene, we calculated the mean of normalised full-length reads (*NFLR*_*Ti*_) across all the samples:

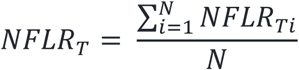

where, *NFLR*_*T*_ represents the mean expression of transcript *T* across all samples and *N* is the total number of samples. To remove low-confidence isoforms arising from artefacts, we only selected isoforms fulfilling the following three criteria: (1) expression of minimum 0.1% of total transcription per sample, i.e., *NFLR*_*Ti*_ ≥ 0.1; (2) a minimum of 80% of total samples passing the *NFLR*_*Ti*_ threshold; and (3) expression of minimum 0.3% of total transcription across samples, i.e., *NFLR*_*T*_ ≥ 0.3.

### Visualizations of transcripts

For any visualization of transcript structures, we have recently developed ggtranscript (*55*) (v 0.99.03; https://github.com/dzhang32/ggtranscript), a R package that extends the incredibly popular tool ggplot2 (*56*) (v 3.3.5 RRID; SCR_014601) for visualizing transcript structure and annotation.

### CAGE-seq analysis

To assess whether predicted 5’ TSSs of novel transcript were in proximity of Cap Analysis Gene Expression (CAGE) peaks we used data from the FANTOM5 dataset (*37*, *38*). CAGE is based on “cap trapping”: capturing capped full-length RNAs and sequencing only the first 20–30 nucleotides from the 5’-end. CAGE peaks were downloaded from the FANTOM5 project (https://fantom.gsc.riken.jp/5/datafiles/reprocessed/hg38_latest/extra/CAGE_peaks/h <u>g38_liftover+new_CAGE_peaks_phase1and2.bed.gz</u>; accessed 20/05/2022).

### CRISPR Cas9 genome editing

The guide RNA (gRNA) was selected from predesigned Alt-R CRISPR-Cas9 gRNAs at Integrated DNA Technologies (https://www.idtdna.com/pages) (Design ID: Hs.Cas9.APP.1.AE. 5’-CGGTAGGGAATCACAAAGTG-3’)

The ribonucleoprotein (RNP) strategy was used for CRISPR/Cas9 gene editing. The preparation of the RNP complex was performed as per the manufacturer’s instructions using the Alt-R™ CRISPR tracrRNA, crRNA and S.p. Cas9 Nuclease 3NLS (IDT). Electroporation was performed (Amaxa 4D, Lonza) for delivery of the RNP complex into iPSCs. Nucleofected cells were seeded into 10cm dishes and expanded in mTeSR1 media + 10 µm ROCK inhibitor (Stem Cell Technologies). Screening of clones was performed via PCR and Sanger sequencing. Indigo software (https://www.gear-genomics.com/indigo/) was used to perform sequence alignment, and to detect single nucleotide variations and insertion-deletion mutations.

Chromosomal stability was confirmed using low coverage whole genome sequencing, whereby reads were mapped to 1000kb bins and processed for mapability and smoothing following published protocols(*57*). Sequencing was performed in collaboration with UCL Genomics.

### Immunocytochemistry

Cells were fixed onto glass coverslips using 4% paraformaldehyde for 15 minutes, and stored at 4°C in phosphate buffered saline (PBS) before immunostaining. Cells were permeabilised by incubating three times in 0.2% triton-X-100 in PBS (PBS-T) for 10 minutes per incubation, before being blocked in 5% bovine serum albumin (BSA) in PBS-T for 30 minutes. Cells were incubated in primary antibodies (Table 1) diluted in blocking solution overnight at 4°C. After three washes in PBS-T, secondary antibodies (Alexafluor) diluted in blocking solution were added for 1 hour in the dark. After a further three washes in PBS-T, 4’,6-diamindino-2-phenylindole (DAPI) was added at 0.1μg/mL for 10 minutes, as a nuclear counterstain. A final three washes in PBS-T were performed before the cells were mounted onto slides using DAKO fluorescence mounting medium. Imaging was performed using Zeiss LSM microscope at 20x magnification and Leica LAS X software.

**Table 1.**
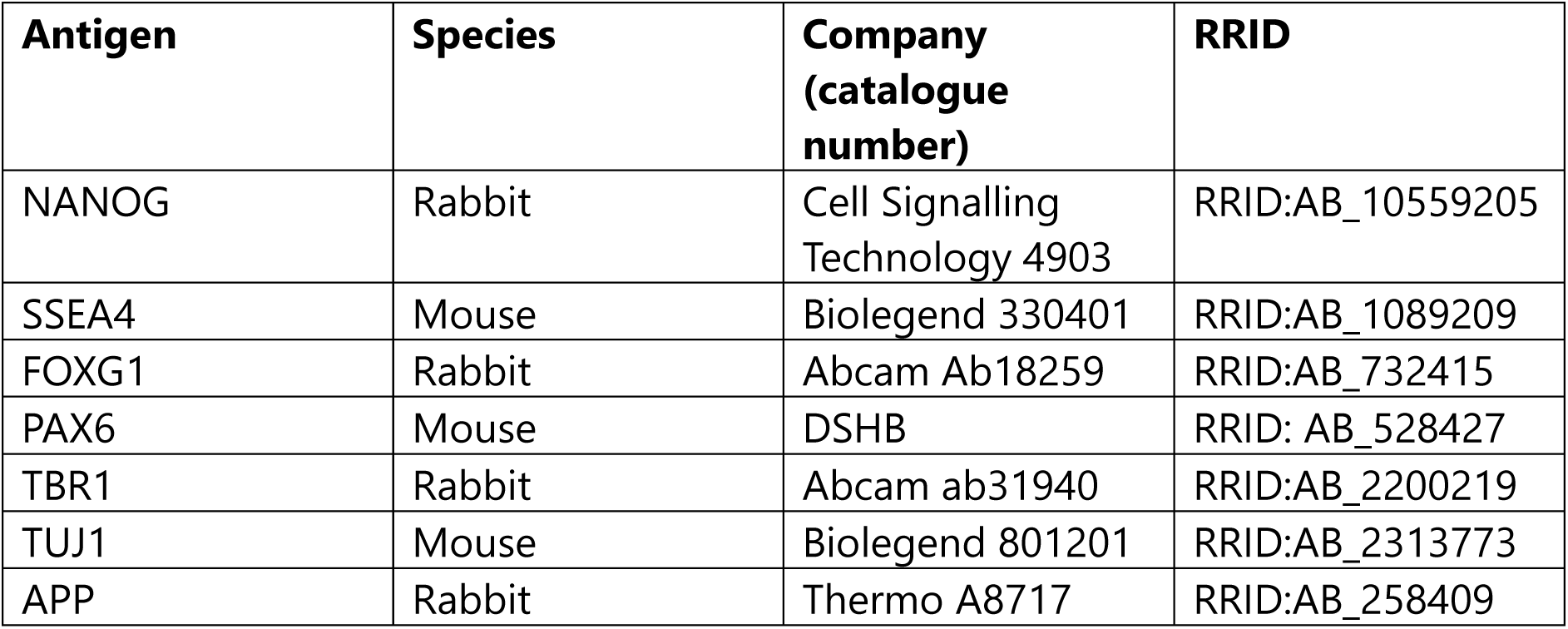
Antibodies for ICC and Western blotting.

### Western blotting

Cells were lysed using RIPA lysis buffer containing protease and phosphatase inhibitors (Roche). Lysates were centrifugated to remove insoluble debris and a BCA assay (BioRad) was performed to quantify protein content. Protein content was standardised across samples, before the samples were loaded with LDS buffer and dithiothreitol (DTT) and denatured by boiling at 95°C for 5 minutes. After centrifugation at 6000 g for 2 minutes at 4°C, proteins were separated via electrophoresis using 4-12% NuPAGE Bis-Tris gel in NuPAGE MES SDS running buffer at 150 V for 1 hour. Transfer onto nitrocellulose membrane was performed at 30 V for 1 hour at 4°C. The membrane was blocked in PBS with 0.1% Tween-20 (PBS-Tween) with 3% BSA. Primary antibodies (Table 1) in blocking solution were added overnight at 4°C. Membranes were washed three times with PBS-Tween and secondary antibodies (Licor Biosciences) in blocking solution were added for 1 hour. A final three washes in 1X PBS were performed before the membrane was imaged using an Odyssey Fc scanner (Licor Biosciences).

### qPCR

cDNA was reverse transcribed from RNA using Superscript IV reverse transcriptase and random hexamer primers (Thermo). qPCR was then performed using POWER Sybr green master mix (Thermo) on an Agilent Aria MX3000 real time PCR machine. For relative quantification, the ΔΔCT method was employed, relative to the housekeeping gene *RPL18A*. RPL18a fwd CCCACAACATGTACCGGGAA, RPL18a rev TCTTGGAGTCGTGGAACTGC, APP fwd GGTACCCACTGATGGTAAT, APP rev GGTAGACTTCTTGGCAATAC, M-C99 fwd ACTAATTGGTTGTCCTGCATACT, M-C99 rev GCCCACCATGAGTCCAATGA.

### Aβ ELISAs

Media was conditioned for 48 hours in neuronal cultures before being collected and centrifuged at 2,000g. Aβ38, Aβ40 and Aβ42 were quantified simultaneously using the Meso Scale Discovery V-Plex Ab peptide panel (6E10), for electrochemiluminescence based quantification. Protein content of each culture was quantified using the BioRad DC BCA assay and this quantification was used to normalise Aβ concentrations to cell numbers.

### Mass spectrometry

Nanoflow liquid chromatography–mass spectrometry analysis of immunoprecipitated CSF samples was performed as described previously (*58–60*). Briefly, to maximise the chance to find low abundant Aβ-peptides, 5 mL CSF was immunodepleted using 3D6 and 2G3 (directed at N-terminal Aβ1 and C-terminal Aβ40, respectively) and then the supernatant was immunoprecipitated using 6E10 and 4G8 (directed at Aβ4-9 and Aβ18-22, respectively) using a KingFisher (Thermo Fisher Scientific) magnetic particle processor (*60*). Immunoprecipitated samples were then analysed using a Q Exactive coupled to Dionex Ultimate 3000 system (both Thermo Fisher Scientific) operating in positive ion mode and with acidic buffers as described in (*60*). Database search was performed using Mascot Deamon v2.6.0 combined with Mascot Distiller v2.6.3 and Mascot database server v2.6.1 for preprocessing and search against a custom made human APP-only database as described previously (*58*, *59*). Fragment ion spectra were validated manually.

## Acknowledgments

We are deeply grateful to the patients who generously donated fibroblasts for this research. We thank all funding agencies for their support. We are also grateful to the UCL Long Read Sequencing Service for their technical assistance. Special thanks to Prof. Dario Alessi, Dr. Pui Yiu Lam, and Dr. Raja S. Nirujogi for their valuable input and guidance throughout this study. For the purpose of open access, the author has applied a Creative Commons Attribution (CC BY) license to all Author Accepted Manuscripts arising from this submission.

## Data and materials availability

All data needed to evaluate the conclusions in the paper are present in the paper and/or the Supplementary Materials. Raw long-read RNA-seq data generated and used in this manuscript are publicly available in Synapse (Project SynID: syn53642785): https://www.synapse.org/#!Synapse:syn53642785/wiki/626685. Other data, code, and materials used in the analysis are available and described throughout the manuscript. The code for analysis and figure generation in this manuscript can be accessed through https://github.com/egustavsson/APP_manuscript.

