## supplementary figures for "Independent Generation of Amyloid-β via Novel *APP* Transcripts"

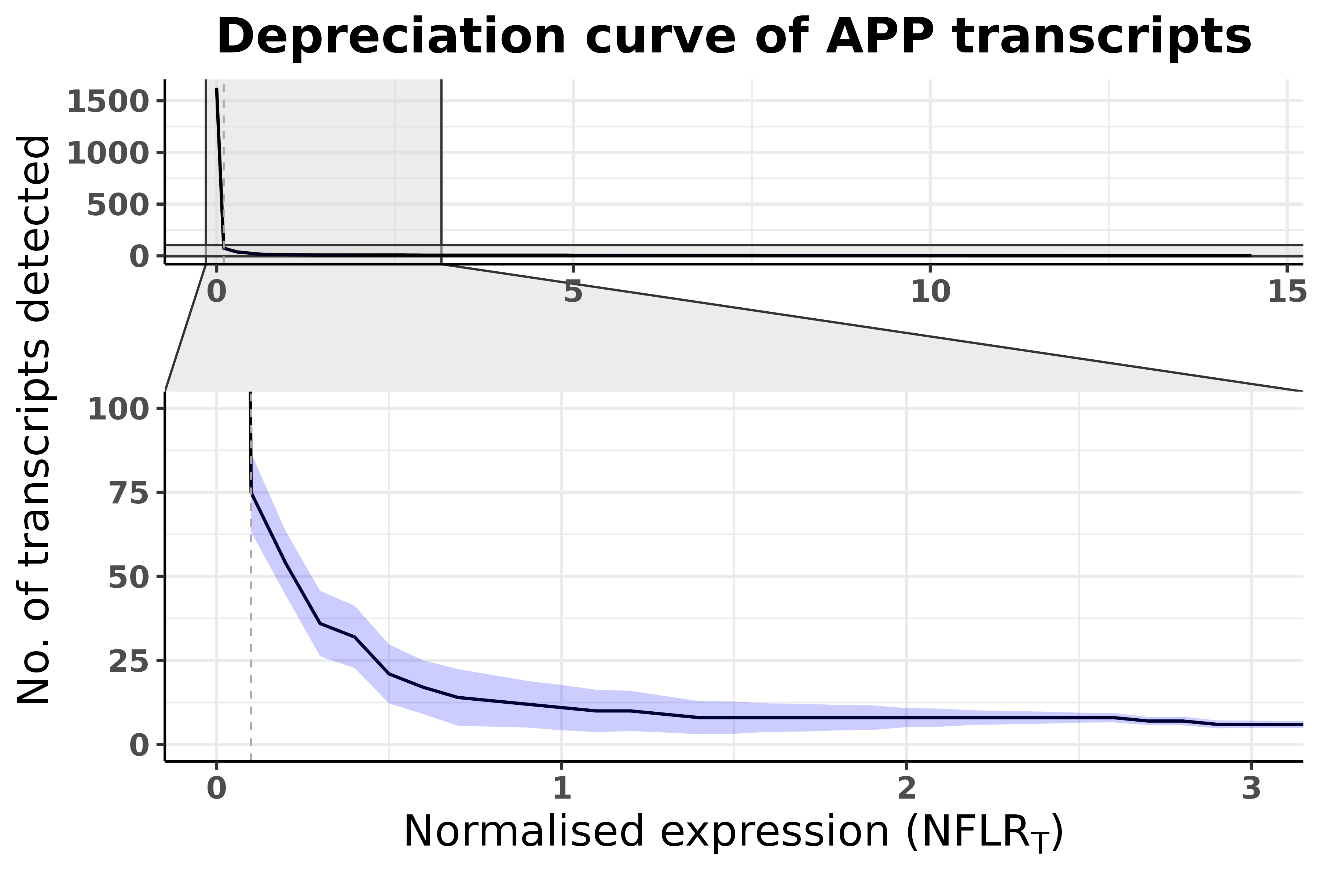

**Supplementary Fig. 1: Total number of unique transcripts of APP by normalized expression.** Depreciation curve showing the number of unique APP transcripts on the Y-axis increased by increasing the normalized full-length read count of transcript (NFLRT) on the X-axis. NFLRT is the total number of reads per transcript normalized by the total number of reads of the loci. The dashed line represent the cut-off used for what is included as a transcript and is set at 0.1, representing a relative expression of 0.1%.

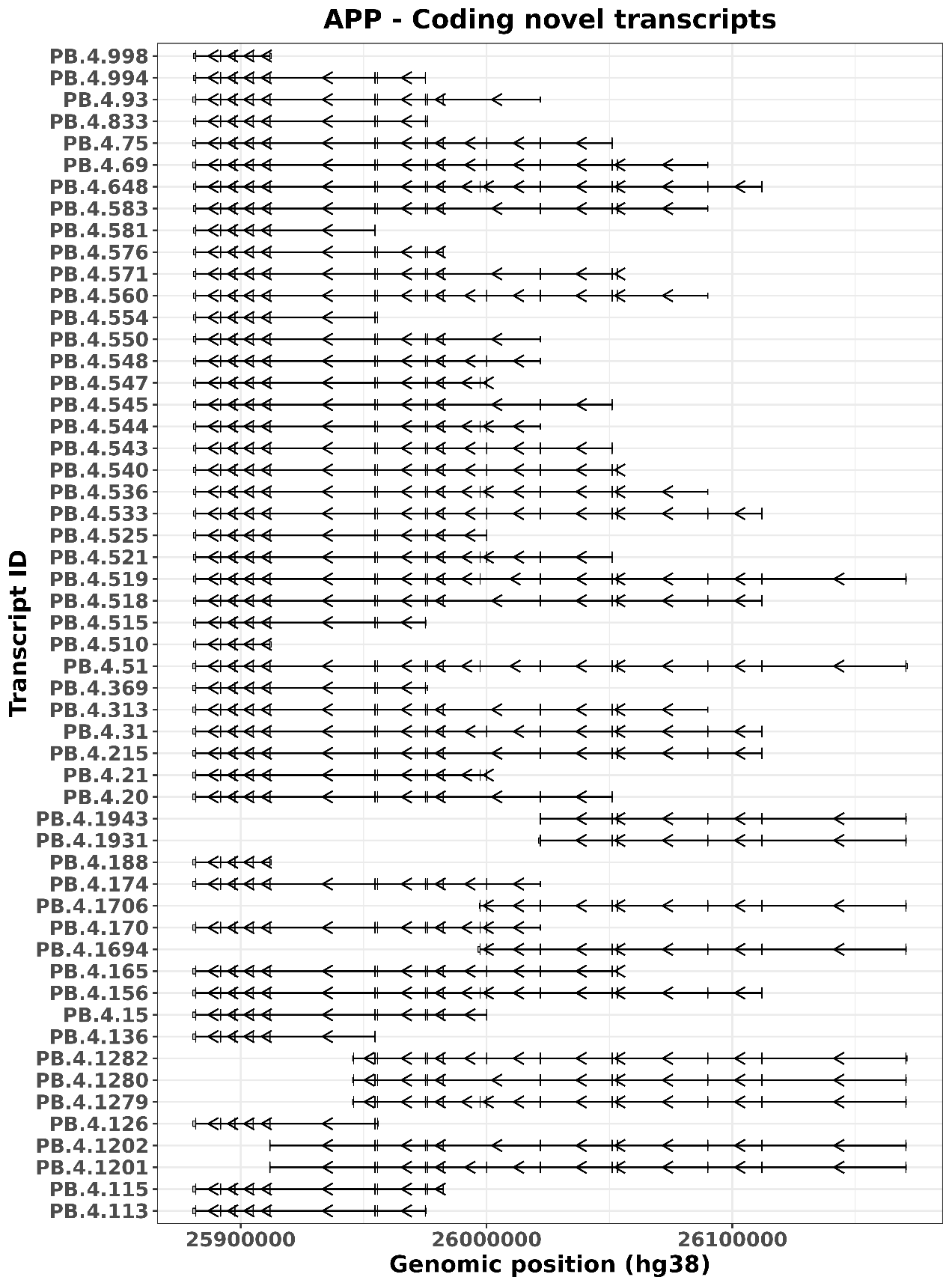

**Supplementary Fig. 2: APP coding novel transcripts.** A total of 54 were predicted to be translated, to avoid nonsense-mediated decay (NMD), and to lack a full-splice match with the reference (classified as coding novel). Transcripts were identified through targeted long-read RNA sequencing across various cell types, including iPSC, neuroepithelial cells, neural progenitor cells, iPSC-derived cortical neurons, astrocytes, microglia, and brain samples.

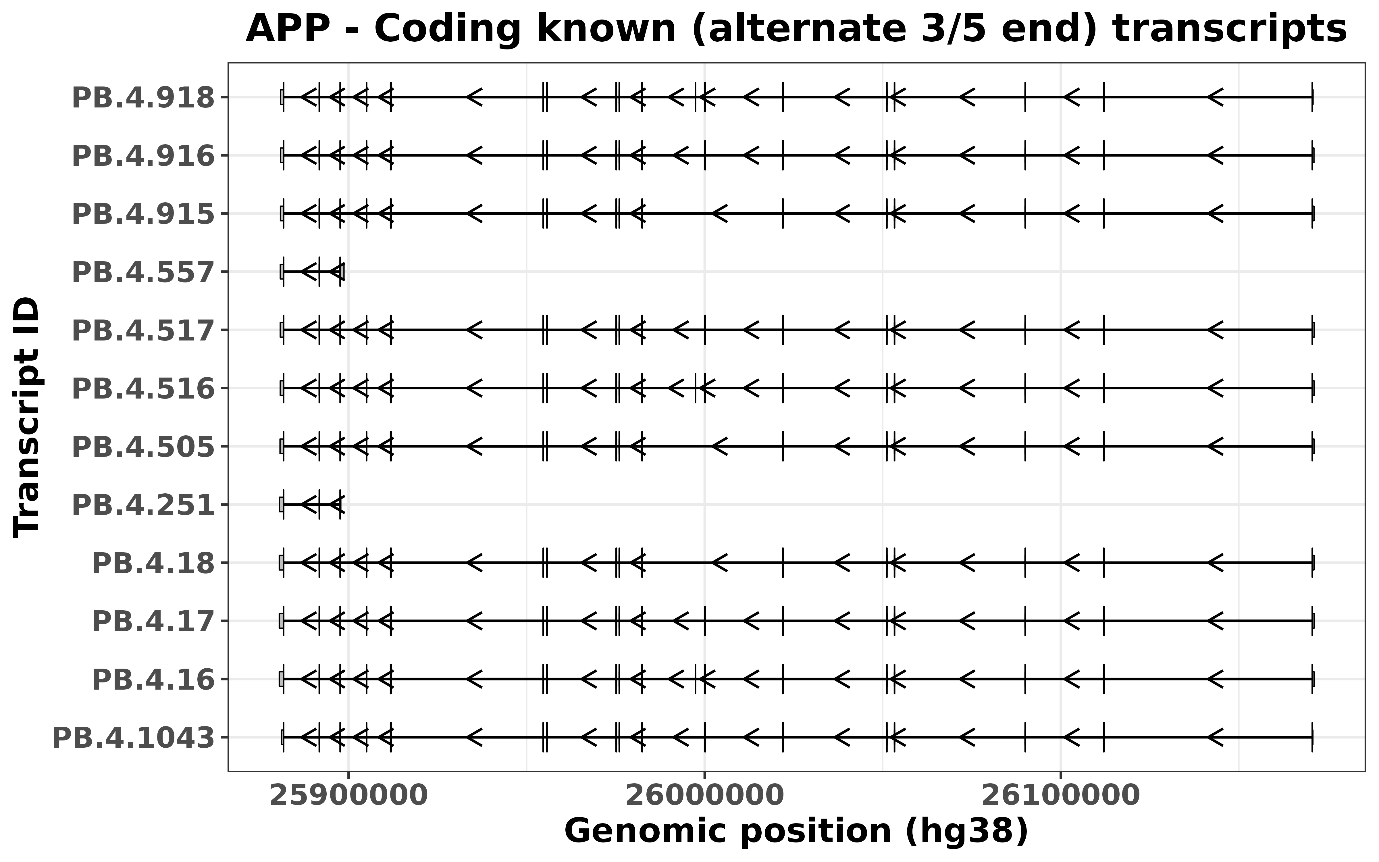

**Supplementary Fig. 3: APP Coding known (alternate 3/5 end) transcripts.** A total of 12 were predicted to be translated, to avoid NMD, and to have a full-splice match with the reference but feature an alternative 3′ end, 5′ end, or both (classified as Coding known (alternate 3/5 end)). Transcripts were identified through targeted long-read RNA sequencing across various cell types, including iPSC, neuroepithelial cells, neural progenitor cells, iPSC-derived cortical neurons, astrocytes, microglia, and brain samples.

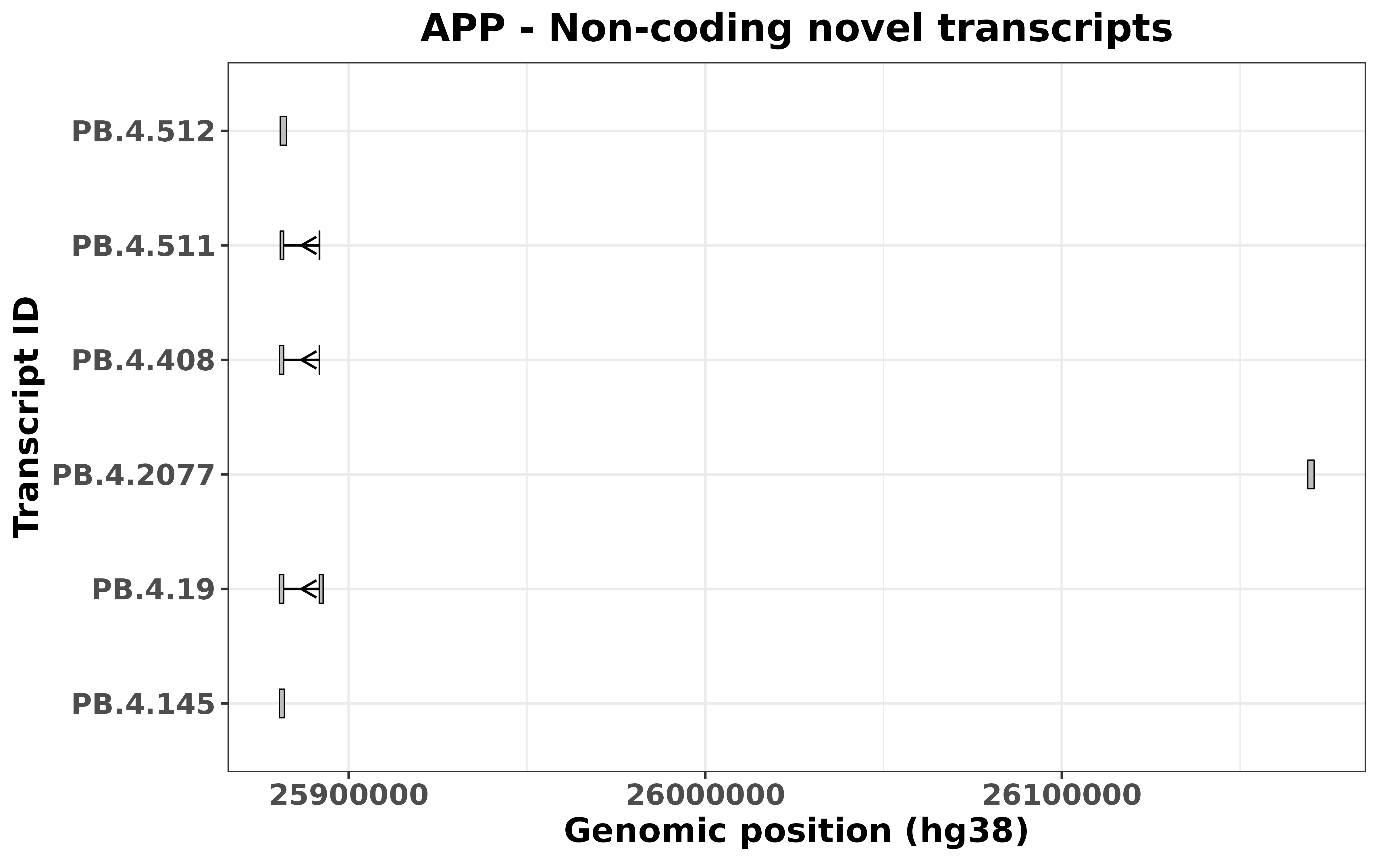

**Supplementary Fig. 4: APP non-coding transcripts.** A total of 4 were predicted to be non-coding and lacked a full-splice match with the reference (classified as non-coding novel). Transcripts were identified through targeted long-read RNA sequencing across various cell types, including iPSC, neuroepithelial cells, neural progenitor cells, iPSC-derived cortical neurons, astrocytes, microglia, and brain samples.

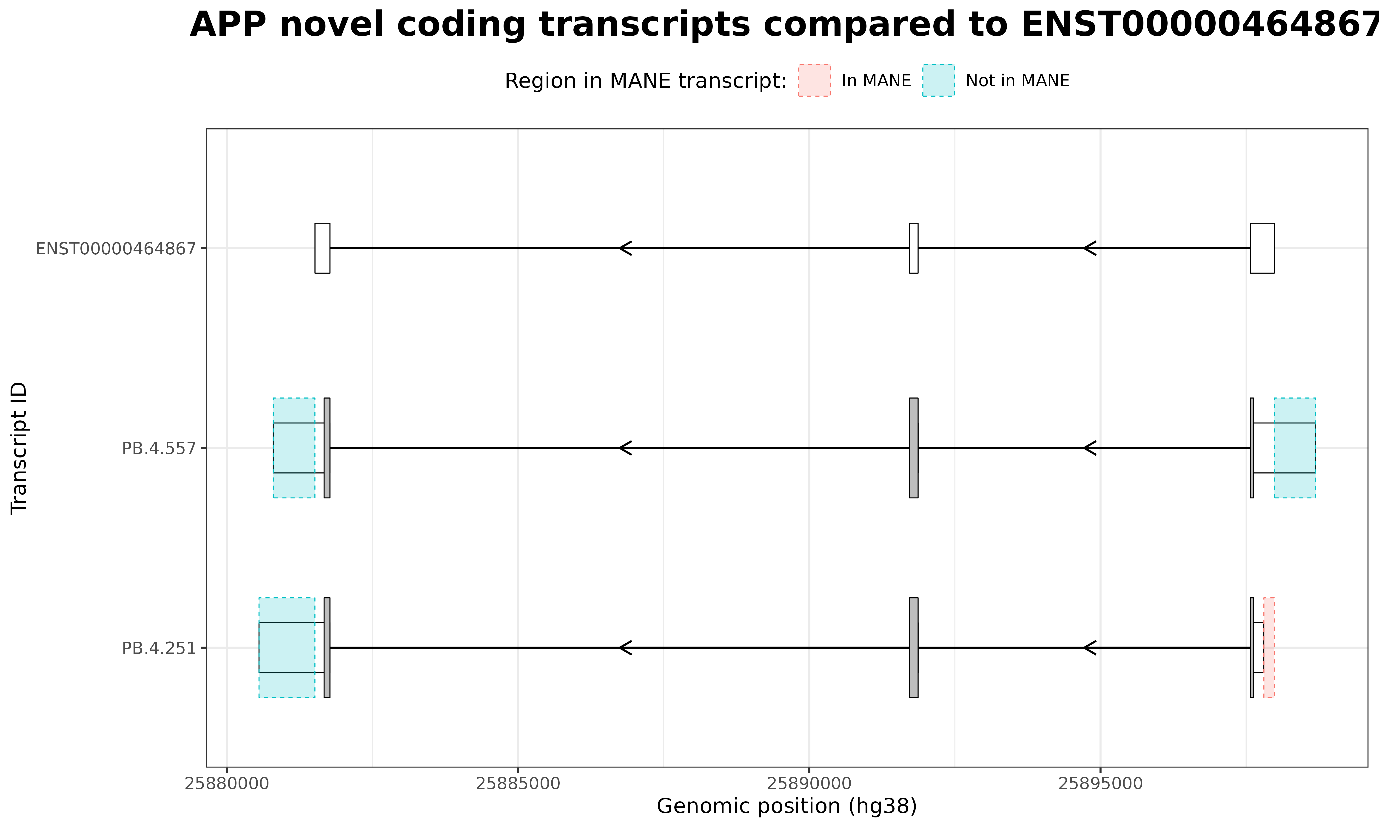

**Supplementary Fig. 4: APP novel coding transcripts compared to ENST00000464867.** Two novel identified transcripts (PB.4.557 and PB.4.251), closely resembles an annotated APP transcript (ENST00000464867) albeit with different lengths of both TSS and TTS at the 5’ and 3’ UTR, respectively.
